# Dynamic compartmentalization in neurons enables branch-specific learning

**DOI:** 10.1101/244772

**Authors:** Willem A.M. Wybo, Benjamin Torben-Nielsen, Marc-Oliver Gewaltig

**Author notes:** Correspondence and requests for materials should be addressed to BTN.

## Abstract

The dendritic trees of neurons play an important role in the information processing in the brain. While it is tacitly assumed that dendrites require independent compartments to perform most of their computational functions, it is still not understood how they compartmentalize into functional subunits. Here we show how these subunits can be deduced from the structural and electrical properties of dendrites. We devised a mathematical formalism that links the dendritic arborization to an impedance-based tree-graph and show how the topology of this tree-graph reveals independent dendritic compartments. This analysis reveals that cooperativity between synapses decreases less than depolarization with increasing electrical separation, and thus that surprisingly few independent subunits coexist on dendritic trees. We nevertheless find that balanced inputs or shunting inhibition can modify this topology and increase the number and size of compartments in a context-dependent, temporal manner. We also find that this dynamic recompartmentalization can enable branch-specific learning of stimulus features.

## Introduction

Brain function emerges from the orchestrated behaviour of billions of individual neurons that transform electrical inputs into action potential (AP) output. This transformation starts on the dendritic tree, where inputs are collected, and proceeds to the axon initial segment where APs are generated that are then transmitted to downstream neurons through the axonal arborization. While axons appear to merely communicate the neuronal output downstream, dendrites collect and non-linearly transform input. This phenomenon is referred to as dendritic computation, and recent experimental work not only demonstrated that it occurs in vivo, but also that it is required for normal brain function (Grienberger et al., 2015; Takahashi et al., 2016; Smith et al., 2013). In both experimental and theoretical work, an abundance of dendritic computations have been proposed (Segev, 2000; Häusser and Mel, 2003; London and Häusser, 2005; Torben-Nielsen and Stiefel, 2010; Silver, 2010). Nearly all of them rely on the tacit assumption that dendrites are compartmentalised into independent subunits: region on the dendritic tree that integrate inputs independently from other regions.

Experimentally, independent compartments have been shown to enable branch-specific control of AP back-propagation (Müllner et al., 2015) as well as local N-methyl D-aspartate (NMDA), Ca2+ or Na+ spikes (Larkum et al., 2007; Wei et al., 2001). These local spikes in turn allow neurons to decode bursts of inputs (Polsky et al., 2009) and they drive the local clustering of correlated synaptic inputs (Lee et al., 2016; Gökçe et al., 2016). Independent compartments also allow plasticity to operate on individual branches (Golding et al., 2002; Losonczy et al., 2008; Govindarajan et al., 2011; Cichon and Gan, 2015; Weber et al., 2016). A recent finding that distal apical dendrites can spike ten-fold more often than somata (Moore et al., 2017) suggests an important role for this branch-specific plasticity (Gambino et al., 2014; Chen et al., 2015; Cichon and Gan, 2015). Independent compartments furthermore allow different input streams to be discriminated from each other (Johenning et al., 2009) and they facilitate sensory perception through feedback signals (Takahashi et al., 2016; Phillips, 2017). Decoupling of spiking in basal dendrites and soma is also implicated in the formation of place fields in hippocampus (Sheffield et al., 2017), while subthreshold independent dendritic activity is thought to guide dendrite growth in auditory neurons (Blackmer et al., 2009). Curiously, recent data has also shown that human brain cells have far more extensive dendritic arborizations than rodent cells (Mohan et al., 2015), leading to the intriguing hypothesis that the number of independent dendritic compartments is related to higher-order intelligence.

Computationally, independent compartments are predicted to enable individual neurons to function as two-layer neural networks (Poirazi et al., 2003b,a). This in turn should enable them to learn linearly non-separable functions, such as XOR (Schiess et al., 2016), and to implement translation invariance (Mel et al., 1998). On the network level, independent compartments are thought to dramatically increase memory capacity (Poirazi and Mel, 2001; Wu and Mel, 2009), to allow for the stable storage of feature associations (Bono and Clopath, 2017), to represent a powerful mechanism for coincidence detection (Larkum et al., 1999; Chua and Morrison, 2016) and to support the back-prop algorithm to train neural networks (Guergiuev et al., 2016).

Because all of these studies rely on compartments in dendrites, one would expect a dendritic compartment to be a well-defined concept. Surprisingly, a rigorous definition of a dendritic compartment does not exist: dendritic compartments have been described as individual branches (Branco and Häusser, 2010), subtrees (Traub et al., 2005), or even small regions of ~ 10 micron (Gökçe et al., 2016; Sheffield et al., 2017). In this work we formalise the concept of a dendritic compartment by linking the dendritic arborization to an impedance-based tree-graph. We show that the topology of this tree-graph reveals independent dendritic compartments and furthermore that the number of compartments can be dynamically regulated in order to perform some of the aforementioned computations.

## Results

### A drastic simplification of the “connectivity matrix” of a neuron

Previous work has established that the eventual compartmentalization of a neuron must be related to its impedance matrix (Cuntz et al., 2010; Zador et al., 1995). In a sense, this matrix can be seen as the “connectivity matrix” of a neuron, as each element *Z_x,x_*_′_ describes how an input current at location x along the dendrite will change the voltage at location *x*′ and hence – through the voltage dependence of the synaptic input currents (see methods) – will influence the processing of inputs at *x*′. Nevertheless, it is not straightforward to determine, from this matrix, exactly where the independent compartments will be. A first complication is that neurons integrate inputs temporally, so that the impedance matrix is in fact a matrix of temporal kernels *Z_x,x_*_′_(*t*) (see methods). A second complication is that the voltage at each location on the dendrite is influenced by the inputs at any other location, both directly, through a contribution *Z_x,x_*_′_(*t*) * *I_x_*_′_(t), and indirectly, through the influence of these terms on the voltage-dependent factors (such as the non-linear unblocking of NMDA receptors (MacDonald and Wojtowicz, 1982; Jahr and Stevens, 1990)) in the local synaptic currents.

To formalize the concept of a dendritic subunit, we first propose a drastic simplification of this impedance matrix, and hence of the underlying dendrite model. We envision the dendritic voltage as a superposition of components that are global, present only in a subtree of the dendrite, or only in local branches (Fig 1A, left). This model can be represented graphically as a tree graph, where each node represents such a voltage component and integrates only inputs arriving into its subtree (Fig 1A, right). We term this tree graph the neural evaluation tree (NET) and illustrate the algorithm we designed to derived it on a granule cell (Fig 1C). The dendritic impedance matrix supports such a tree-like description: it can be seen that a relatively even blue surface covers most of the matrix (Fig 1D). This represents transfer impedances between main dendritic branches. Closer to the diagonal, squares of light blue are present, representing dendritic subtrees whose branches are electrotonically closer to each other than to other branches. On the diagonal, one finds small squares of green and red colors, representing individual terminal branches. We designed a recursive algorithm (see methods for an in-depth description) that associates a temporal kernel with each of these regions: first, a global kernel is defined as the average of all impedance kernels that relate inputs on different main branches (Fig 1E). Then, regions of the impedance matrix that are electrotonically closer to each other than to other regions are identified by determining where the input impedance (the diagonal of the impedance matrix) is in the same range as the global impedance (dashed blue line in Fig 1F). Connected regions outside of this range then form the child nodes of the root node (*c*_0_ to *c*_5_ in Fig 1F), and the procedure is repeated for each child node with the impedance matrix restricted to regions associated with the child node (red squares in Fig 1G). Crucial for our simplification is the observation that impedance kernels with equal steady-state impedance values (i.e. the integral over the kernel) share similar time-scales (Fig 1B), so that averaging them into a single kernel indeed yields accurate dynamics.

**Figure 1:**
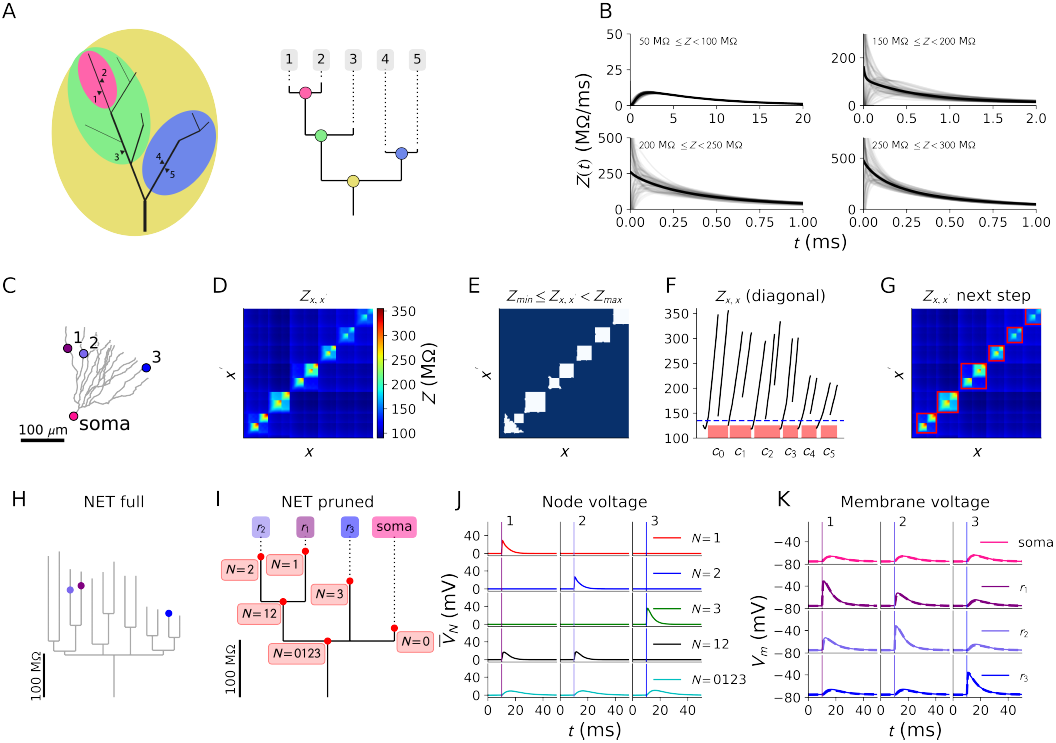
Construction of the NET. **A:** Cartoon of the idea behind the NET: the dendrite voltage is obtained as a superposition of voltage components (left) that are either global (yellow), span a dendritic subtree (green, blue) or a single branch (pink). The structure of this superposition can be represented graphically as a tree graph (right). **B:** The average impedance kernels for impedances within the indicated ranges on a granule cell. **C:** The granule cell dendrite with three inputs regions (numbers 1–3) and somatic readout. **D:** The impedance matrix associated with the granule cell. Color encodes the impedance between each pair of points. **E:** Points in the impedance matrix of the granule cell where the impedance is between *Z*_min,0_ (i.e. the minimal transfer impedance in the cell) and *Z*_max,0_(= 135 MΩ, chosen for illustrative purposes) are coloured blue. The average of the impedance kernels associated with these points will constitute the impedance kernel of the NET root. **F:** The input impedance (diagonal of the impedance matrix). Connected domains with input impedance larger than *Z*_max,0_ are indicated on the x-axis and denoted by *c_i_*, as they will constitute the child nodes of the root in the NET. **G:** Dendritic domains associated with each child node *c_i_* are marked by a red square on the impedance matrix. These restricted matrices are used in the next step of the recursive algorithm. **H:** The full NET for the granule cell. Nodes where inputs from regions 1 to 3 arrive are indicated in the corresponding color. **I:** The pruned NET obtained by removing all nodes that do not integrate regions 1 to 3. **J:** Voltages at each node in the NET are computed. Leaf nodes (*N* =0, 1, 2, 3) contain voltage components unique to the associated regions while lower nodes (*N* = 12 and *N* = 0123) contain voltage components that are shared between regions. Voltages contained in each node are shown after an input is presented in location 1 (left), 2 (middle) and 3 (right). Onset of the input is indicated with a vertical line in a color matching with the input location in (C). **K:** Final voltages calculated for the soma and input regions (*r*_1_ -*r*_3_) by summing the node voltages on the path from root to associated leaf. Dashed lines are computed using the NET and solid lines are the voltages from simulating an equivalent NEURON model.

To study interactions between a subset of input regions on the dendritic tree, the full NET is pruned (see methods) so that only nodes that integrate the regions of interest are retained (Fig 1H, I). It is instructive to illustrate how the dendritic voltage at these regions of interest is constructed by the NET: each nodal voltage component is computed by convolving the impedance kernel at that node with all inputs in its subtree (Fig 1J). The local dendritic voltage at a location is then constructed by summing the NET voltage components of all nodes on the path from root to leaf (Fig 1K), so that the local voltage is indeed a superposition of both global and local components. Note that this local voltage coincides with the voltage computed through simulating an equivalent NEURON model (Carnevale and Hines, 2006) (dashed versus full line in Fig 1K). The key feature here is that the voltage components of the NET leafs (nodes *N* = 1, 2, 3) feature no direct dependence on inputs at other locations (Fig 1J), in contrast to the local voltages in the biophysical model (Fig 1K). Note furthermore that while we have only implemented AMPA synapses in this example, it is straightforward to implement other voltage dependent currents, such as NMDA synapses or voltage-gated ion channels, in the NET framework (see methods).

In conclusion, by formulating the NET framework to model dendritic trees, we have markedly simplified the task of finding independent compartments in two ways: we have strongly reduced the spatial and temporal complexity of the impedance matrix while at the same time introducing voltage components (at the NET leafs) that only depend on local inputs. The only contribution that remains to be quantified is the influence of global NET components on the voltage dependent factors in the local synaptic currents. After validating the NET framework, we will turn our attention to this question.

### Validation of the NET framework

To validate the NET framework, we derived NETs for three exemplar morphologies (Wang et al., 2002; Carim-Todd et al., 2009; Hay et al., 2011) (Fig 2A-C). We then compared the exact impedance matrix with its NET approximation (Fig 2A-C, top right panels). The small value of the root mean square error (RMSE) between both matrices suggests that the NET will accurately reproduce the full neuronal dynamics (Fig 2D). To ascertain this, we equipped each NET with somatic AP channels and simulated it for 100 s while providing Poisson inputs to 100 randomly distributed excitatory and inhibitory synapses (see methods for all the simulation parameters). The resulting voltage traces coincide with the traces obtained from equivalent NEURON simulations (Carnevale and Hines, 2004) (Fig 2A-C, bottom right panels). Consequently somatic RMSEs are low while spike prediction is excellent (Fig 2E).

**Figure 2:**
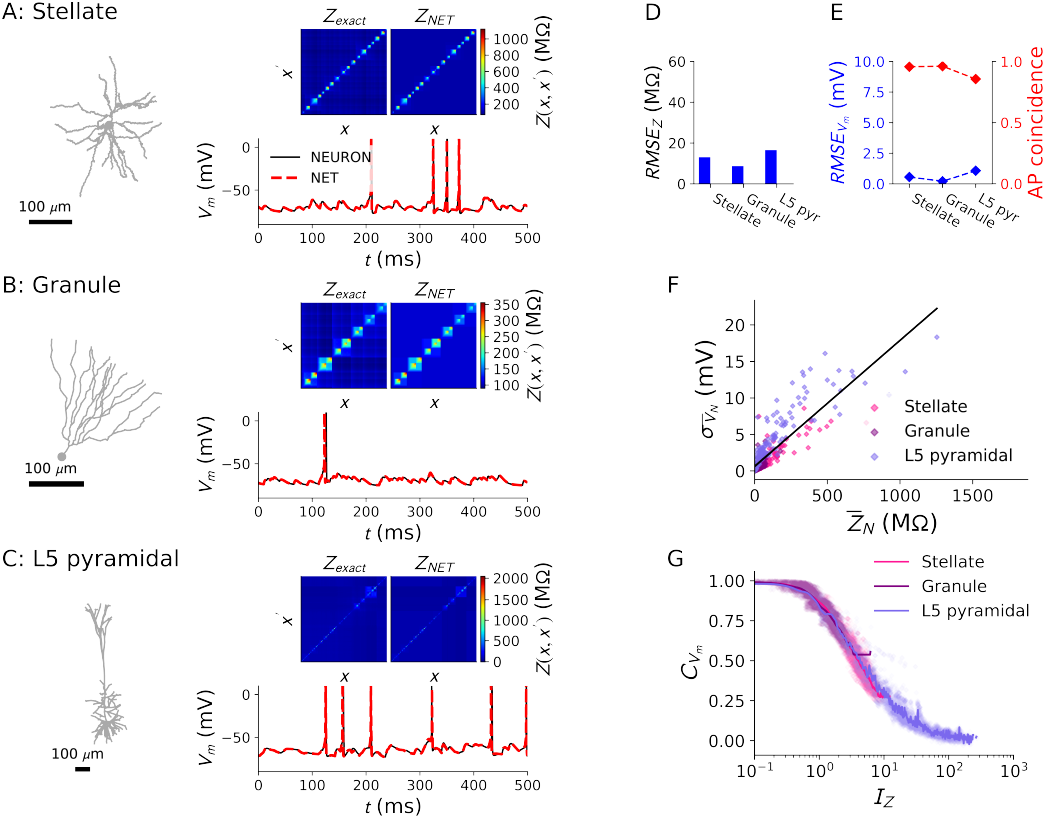
Validation of the NET framework and the impedance-based independence index *I_Z_*. **A:** Cortical stellate cell. **B:** Hippocampal granule cell. **C:** Cortical L5 pyramidal cell. For each cell, morphology is shown on the left, the exact impedance matrix and its NET approximation on the top right and the somatic voltage traces obtained from equivalent NET and NEURON simulations on the bottom right. **D:** The RMSE between NET and exact impedance matrices for each cell. **E:** The subthreshold somatic voltage RMSE (blue) and the spike coincidence factor (red). **F:** The size of the fluctuations of each nodal voltage component (quantified by its standard deviation) as a function of the associated nodal impedance. The black line is a least-squares fit to the data. **G:** The voltage correlation between each pair of synapses on the three cells as a function of *I_Z_*.

### A single number that describes the degree of independence between regions

In the previous paragraphs we have established that NET leafs only receive local inputs. These leaf voltage components hence only integrate local synaptic input currents (modelled as the product between synaptic conductance and voltage dependent factors *I*_syn_(*t*) ≡ *g*_syn_(*t*) *f* (*V*(*t*)), see methods for details). As per the NET framework, the local voltage can be decomposed as

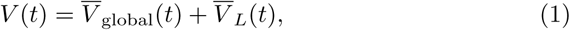

where 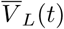 is the voltage associated with the leaf node and 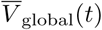is the sum of all other voltage components on the path from root to leaf (note that we denote NET-related quantities with a line over the variable). The leaf voltage is computed by convolving the synaptic input current with the impedance kernel 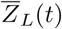 associated with the leaf node:

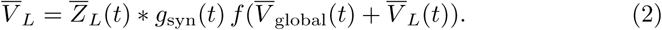

From this equation, it can be seen that 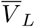 will be independent from all other synaptic inputs if the fluctuations in 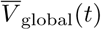 are small compared to the fluctuations in 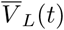. Conversely, if the fluctuations in 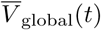 are sizeable compared to the fluctuations in 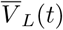, there will be a large degree of cooperativity between the region that leaf *L* integrates and other dendritic regions.

We observe that the voltage fluctuations at each node are approximately proportional to the associated nodal impedances (Fig 2F). This led us to hypothesize that independence between pairs of input regions would be well predicted by the ratio of local over shared impedance:

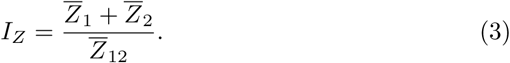

We term this ratio the “impedance-based independence index.” We plotted the membrane voltage correlations between each pair of synapses on our exemplar cells, obtained from our validation simulations (Fig 2A-C), as a function of *I_Z_*. These correlations decrease uniformly as a function of *I_Z_* (Fig 2G). Moreover, the functional dependence of this decrease on *I_Z_* is the same for each cell, indicating that compartmentalization is well predicted by *I_Z_* and other morphological features only play a minor role (see also Fig S1F-H). Note that we visualize the degree of independence between dendritic regions by plotting the vertical lengths of the branches leading up to NET nodes proportional to their associated impedance (Fig 1I).

### The threshold for full independence

As can be seen from Fig 2G, independence between regions on the dendrite increases in a continuous fashion as a function of *I_Z_*. We nevertheless wondered whether there is a threshold value for *I_Z_* above which regions constitute truly independent subunits. Experimentally, the idea has emerged that every branch is a dendritic subunit (Branco and Häusser, 2010). Sharp drops in transfer impedance across bifurcation points lead to this idea, as delivering relatively few stimuli to a single dendritic branch, for instance through two-photon glutamate uncaging (Pettit et al., 1997), leads to a marked attenuation of the measured signal – typically Ca2+ dye luminescence – across bifurcation points (Wei et al., 2001) (Fig 3A). Nevertheless, inferring the extent of the dendritic subunits from this signal is problematic in two ways. First, the Ca2+ signal itself is thought to be more local than the voltage signal (Simon and Llinás, 1985; Biess et al., 2011) while also being influenced by the non-linear activation function of voltage-dependent calcium channels (Almog and Korngreen, 2009). Second, even though there may be a relatively large amount of voltage attenuation across bifurcation points (Fig 3B, C), the remaining depolarization may still be sufficient to cause a large degree of cooperativity between synapses during an ongoing barrage of inputs. To test this idea, we constructed a toy model with two leafs and one root representing, for instance, a single trunk that bifurcates into two child branches, that allowed us to vary *I_Z_* on a log scale between .1 and 100 (Fig 3D, left panel). We then stimulated synapse 1 with a supra-threshold conductance (strong enough to elicit an NMDA spike) and measured the depolarization in branch 1 and in branch 2. It can be seen that there is substantial voltage attenuation: at *I_Z_* = 1 only 50% of the depolarization in branch 1 arrives at branch 2 (Fig 3D, right panel). Nevertheless, membrane voltage correlations decrease much more slowly as a function of *I_Z_*, so that for *I_Z_* = 1 this correlation is still close to 1 (Fig 2G).

**Figure 3:**
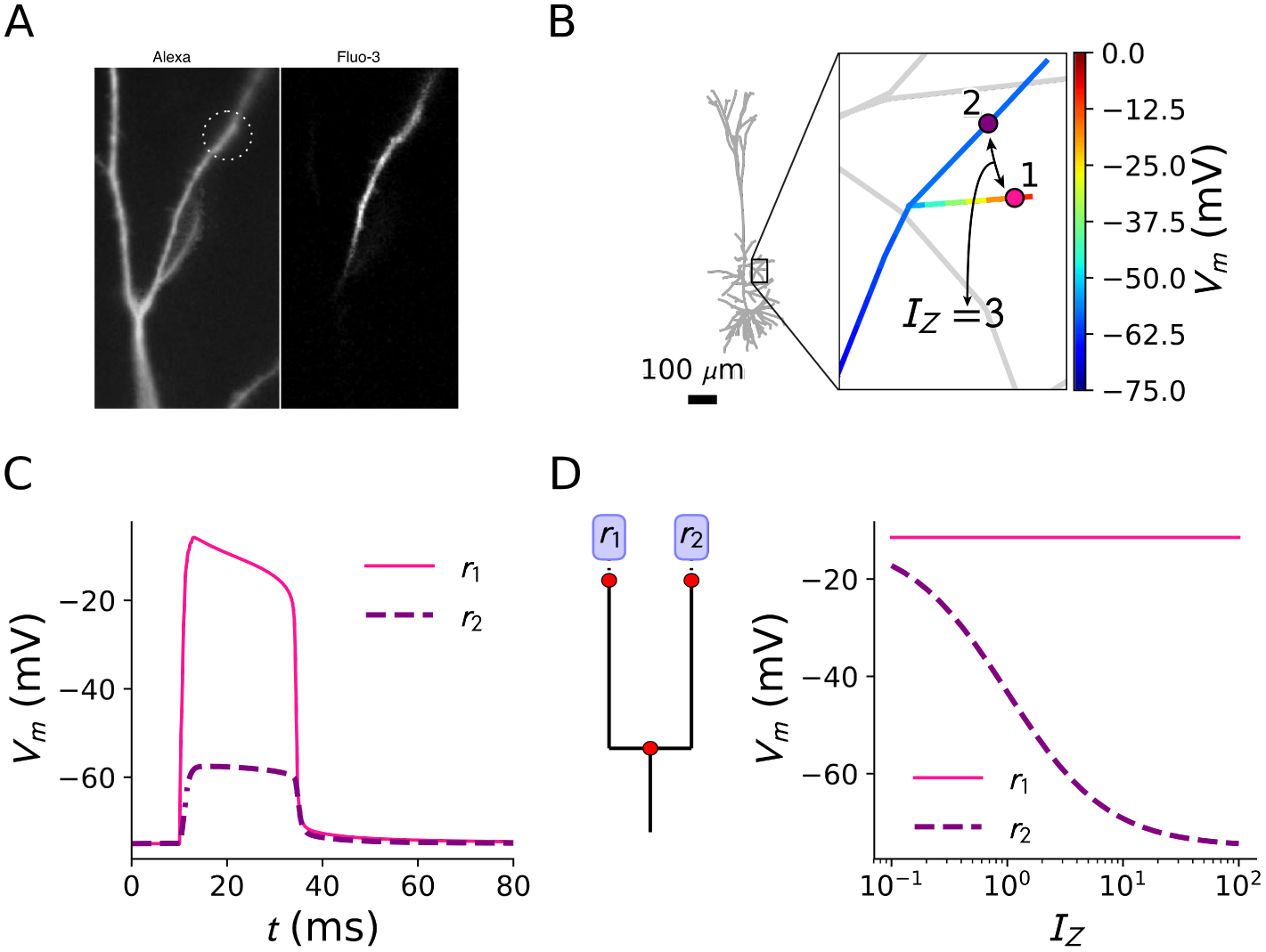
Voltage attenuation makes dendritic branches look highly compartmentalized. **A:** Example of a typical experimental situatio**n:** a branch is stimulated through two-photon glutamate uncaging (dashed circle, left panel) and a correlate of the voltage (here Ca2+, right panel) is measured (adapted with permission from (Wei et al., 2001)). **B:** Voltage distribution (computed with a neuron simulation) in an oblique apical fork 10 ms after supra-threshold stimulation of an NMDA synapse at region 1. A rapid decrease of the depolarization makes the branches seem highly compartmentalized. *I_Z_* =3 between regions 1 and 2. **C:** Trace of the depolarization at regions 1 and 2. **D:** Toy NET model, representing a dendritic fork, where *I_Z_* could be varied at will (left). Depolarization at regions 1 and 2 is shown as a function of *I_Z_* following an input to region 1 (right).

We thus aim at constructing a criterion for independence between dendritic regions that is valid when the dendritic tree receives an ongoing barrage of input. To do so, we construct a control NET where the leaves are completely independent by replacing shared nodal voltage components (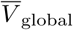 in equation (2)) in the synaptic currents by their long term average (here computed as a 200 ms low-pass filter of the shared voltage under Poisson stimulation with a fixed rate). Consequently, in this independent NET (iNET) leaf voltage components – and thus the synaptic currents – only depend on the local inputs by construction. Convergence between the iNET and the normal NET then indicates independence between input regions. We illustrate the iNET in a toy model with two leafs and one root (Fig 4A). When a synaptic input arrives at synapse 1, an NMDA event will be generated in branch 1. The leaf voltage associated with this event is shown in the leftmost panels in Fig 4B. The effect of a strong input at synapse 2 on the dynamics in branch 1 depends on *I_Z_* : for *I_Z_* = 3, the shape of the NET voltage trace at leaf 1 is drastically modified, whereas at *I_Z_* = 10, there is barely any change (Fig 4B, rightmost panels). In the iNET, the traces at leaf 1 with or without input to synapse 2 are identical by construction. Hence, when the dynamics of the NET converge to those of the iNET, the associated input regions formally constitute independent subunits. We stimulated both input sites with Poisson inputs, and at *I_Z_* = 3 the iNET voltage traces at soma and leafs deviate significantly from their NET counterparts (Fig 4C, top panels). The cooperative dynamics between input sites are thus non-negligible. At *I_Z_* = 10 the iNET and NET dynamics agree very well (Fig 4C, bottom panels). Analysing the voltage correlations between both leafs confirms this finding: for *I_Z_* ≳ 10 the NET correlation approaches the iNET value of zero (Fig 4D). The RMSE between the somatic traces also vanishes then. These results generalize to realistic neuron models and multiple input regions: analysis of leaf voltage correlations and somatic RMSEs for NETs obtained by distributing four input regions on the pyramidal cell morphology (Fig 2C) yields similar results (Fig 4E, the x-axis represents the average between the input regions). While independence increases continuously as a function of *I_Z_*, we conclude, as a rule of the thumb, that for *I_Z_* ≳ 10 pairs of dendritic regions can be considered independent. A further set of criteria concerning the interaction of local non-linearities with each other and with plasticity leads to the same conclusion (Fig S1A-E).

**Figure 4:**
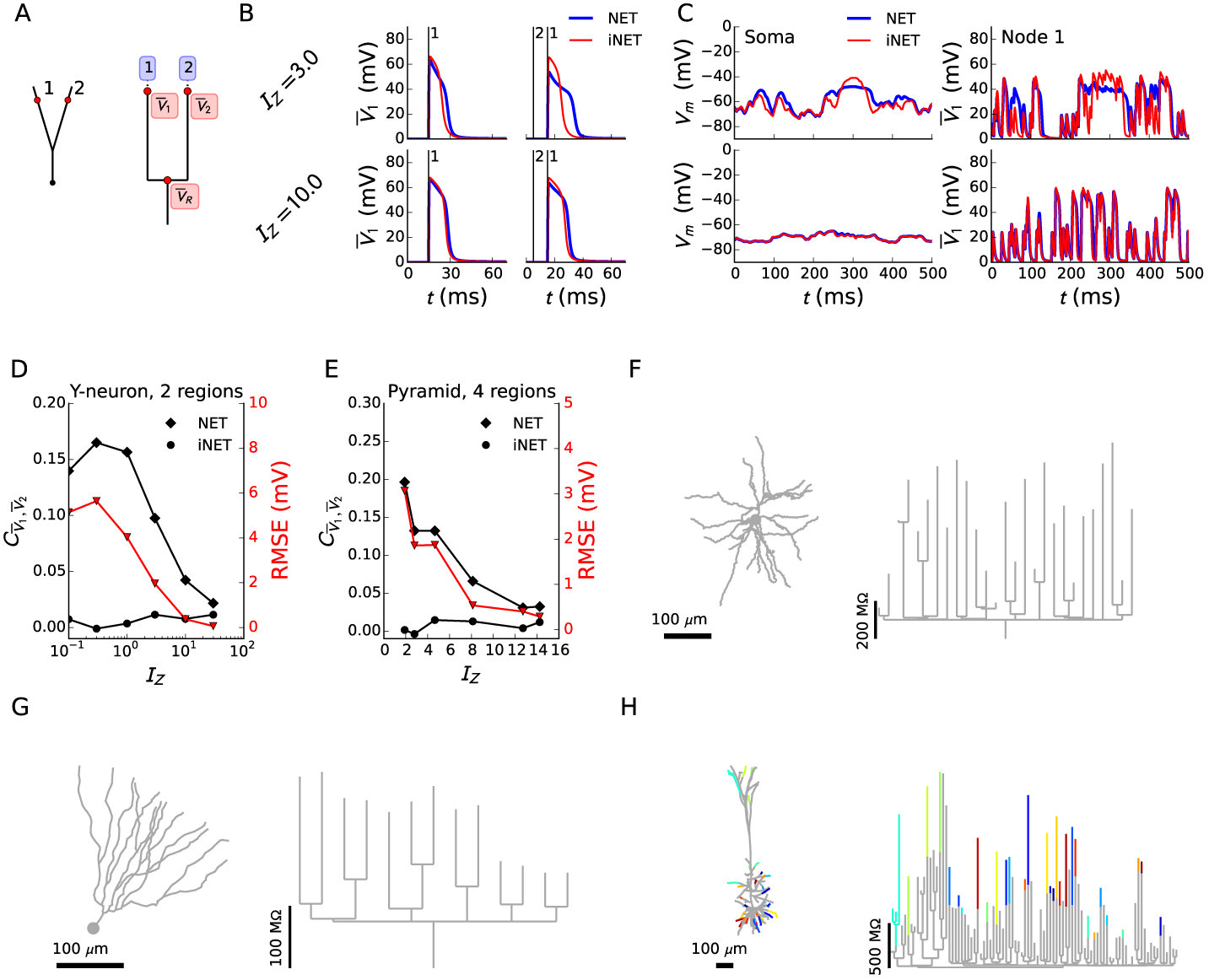
The impedance-based independence index *I_Z_* leads to a systematic characterization of independent compartments. **A:** Toy model morphology and associated NET. **B:** Voltage associated with leaf node 1 for *I_Z_* = 3 (top) and *I_Z_* = 10 (bottom). The iNET trace (red) is compared to the NET trace (blue). In the left column, one input is delivered to synapse 1. In the right column, an input to synapse 2 precedes the input to synapse 1 (vertical black lines indicate input arrival). **C:** Same model, but now both synapses receive Poisson inputs. Somatic voltage (here equal to 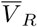, left) and node voltage (right) are shown for *I_Z_* = 3 (top) and *I_Z_* = 10 (bottom). **D:** The correlation between voltages in the leafs of NET and iNET (black), and their somatic RMSE (red), as a function of *I_Z_*. **E:** Same as in D, but with each time 4 inputs regions selected on the L5 pyramidal cell. The x-axis represents the average *I_Z_* between regions. **F:** Cortical stellate cell. **G:** Hippocampal granule cell. **H:** Cortical L5 pyramidal cell. For each cell, the morphology (left) with color coded independent regions at *I_Z_* = 10 and the NET with the same color coded independent compartments (right) are shown. Note that individual compartments are always connected, and color matches between separated compartments are purely accidental.

### A formal definition of dendritic compartmentalization

Next, we asked how many independent regions could maximally coexist along a dendritic tree, as well as what their location would be. Using the NET, we constructed an algorithm that divides the dendritic tree into regions separated by a minimal *I_Z_* (see methods). The resulting compartmentalization is shown in Fig 4F-H. For each exemplar cell, we show the original dendrite morphology (left) as well as the NET (right) with the compartments color-coded. Note that in the pyramidal cell, the number of compartments for *I_Z_* ≥ 10 was far less than the number of dendritic terminals (Fig 4H), and in the stellate and granule cells we did not identify any compartments (Fig 4F, G). Even at lower *I_Z_* values, there are still much less independent compartments than dendritic terminals (Fig S2, see for instance compartment numbers for *I_Z_* ≥ 3). We conclude that, because cooperativity between synapses decreases much more slowly as a function of the electrical separation than voltage attenuation, the amount of independent subunits that can coexist on a dendritic tree is surprisingly low. Our results thus contradict the notion that every branch constitutes an independent *electrical* subunit.

### Dendritic compartmentalization can be modified dynamically by inputs

As neurons perform different input-output transformations at different moments in time, such as during up/down states (Wilson and Kawaguchi, 1996) or attention (Buschman and Kastner, 2015), we hypothesized that the number of compartments in dendrites can be modified dynamically by spatio-temporal input patterns. In principle, shunting inputs could mediate such a dynamic change. Indeed, the voltage obtained at a location with impedance kernel *Z*(*t*), a shunting conductance *g*_shunt_ and excitatory conductances *g_e_*:

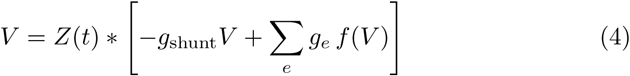

can be reinterpreted in the following way:

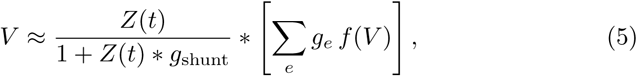

so that the shunt effectively reduces the impedance. When shunts are placed on the dendrite in such a way that they primarily affect shared nodes, independence between branches would increase.

In the high-conductance state (Destexhe et al., 2003) – a state occurring in vivo when many synapses are randomly activated – the conductance of the membrane is increased over the whole dendritic tree, implementing a global shunt. A priori, two things could happen: either the root impedance of the associated NET could decrease more than the leaf impedances, in which case the number of subunits would increase, or the root impedance could decrease less than the leaf impedances, in which case the number of subunits would decrease. Because the root node of the associated NET integrates much more inputs than more local nodes, the former tends to decrease more than the latter (Fig 5B). As a result, electrical separation becomes stronger between branches that only have the root in common, and compartments emerge in the stellate cell (Fig 5D). The high conductance NET is accurate in reproducing the average voltages: traces in Fig 5C (full coloured lines) agree very well with the average post-synaptic potentials computed during an ongoing barrage of balanced excitation and inhibition (dashed coloured lines).

**Figure 5:**
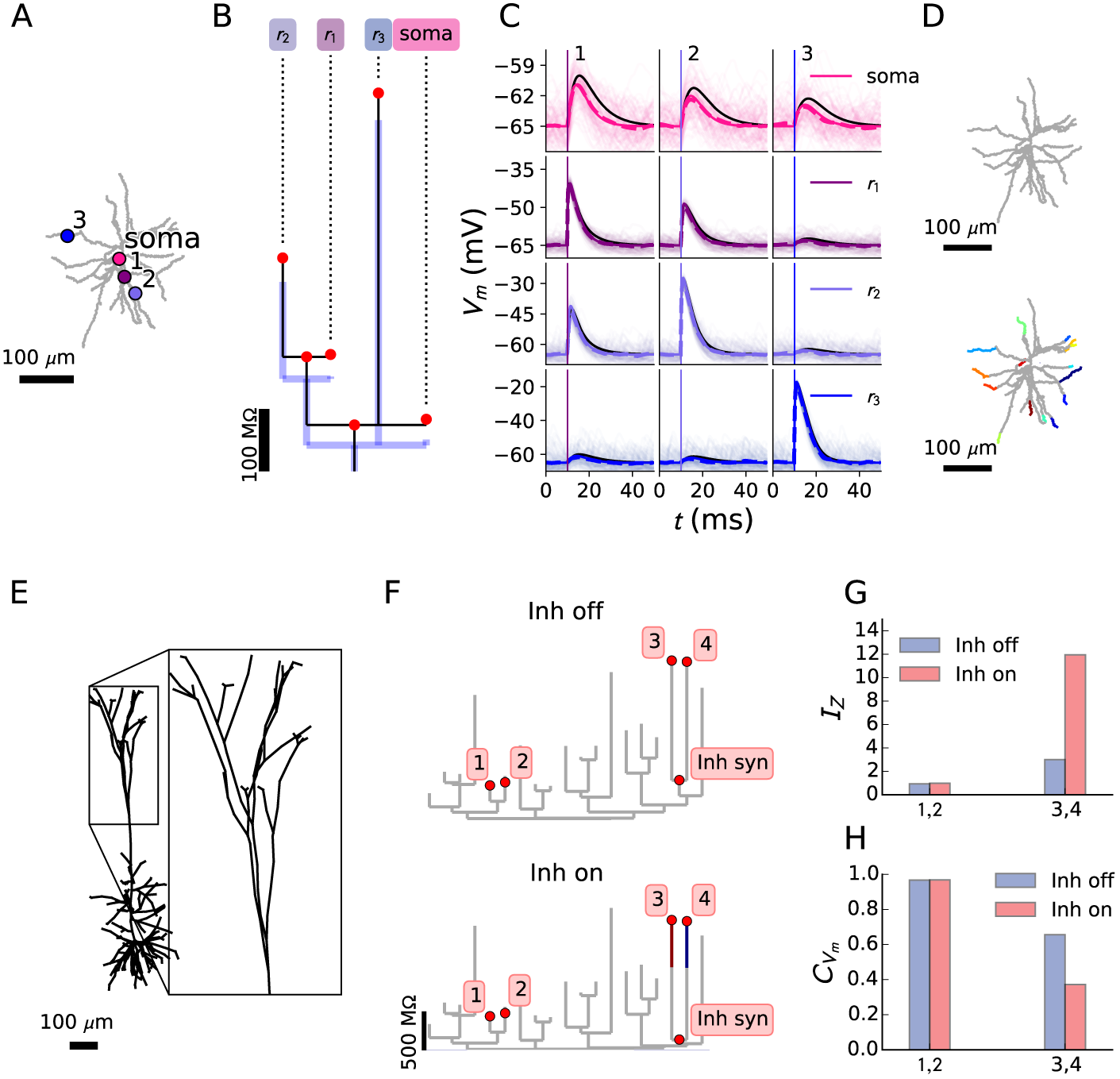
Dynamic compartmentalization due to spatio-temporal input patterns. **A:** Stellate cell morphology with three input regions (numbers 1–3) and somatic readout. **B:** NET for the configuration in A. To mimic the high-conductance state, 200 excitatory and inhibitory synapses were distributed randomly on the neuron and activated with Poisson spike-trains. The NET was recomputed with the time-averaged synaptic conductances as static shunts (blue). **C:** At the three regions of interest, a strong excitatory synapse was inserted. The average post-synaptic potential was computed over 100 trials (dashed line) and coincides with the NET prediction (full line). The responses without background activity are plotted in black for reference. **D:** Compartmentalization for *I_Z_* ≥ 10 in the rest state (top) vs. the high conductance state (bottom). **E:** The effect of inhibition on compartmentalization in the apical tuft of the L5 pyramidal cell was studied. **F:** NET associated with the apical tuft, without (top) and with (bottom) inhibition (with an time-averaged conductance of 5 nS). **G:** *I_Z_* change when inhibition is turned on and **H:** associated change in membrane correlation when synapses in both branches were stimulated with random Poisson trains.

Since both theory (Gidon and Segev, 2012) and connectivity data (Bloss et al., 2016) suggest the importance of the precise location of synaptic inhibition, we investigate the influence of precisely located inhibitory inputs on compartmentalization. In the apical tree of the L5 pyramidal neuron (Fig 5E) we noticed that the sibling branches in a particular apical fork did not constitute separate compartments (locations 3 and 4 in Fig 5F). Upon inserting inhibitory synapses near the branching point between the two terminal segments, they separate into independent compartments (Fig 5F, bottom panel). The required inhibition (with a time-averaged conductance of 5 nS) could be provided by the somatostatin-positive interneuron pathways targeting the apical tuft (Munoz et al., 2017; Markram et al., 2015). This change in compartmentalization can be quantified by the change in *I_Z_* (Fig 5G). Note that the effect is location-specific: independence between locations 1 and 2 does not increase. These predictions are confirmed by computing the voltage correlation between both pairs of branches under Poisson stimulation (Fig 5H).

### Dynamic compartmentalization can enable branch-specific learning

Recent experiments have demonstrated that inhibitory interneurons are required for branch-specific plasticity (Cichon and Gan, 2015). Can a transient recompartmentalization, mediated by inhibition, underlie this branch-specific learning? Before learning, post-synaptic targeting is thought to be unspecific (Gerstner et al., 1996). Hence, inputs coding different stimulus features can arrive at the same branch, but with different strengths. We ask whether sibling branches can learn to become selective only to the strongest initial feature (Fig 6A), using only NMDA spikes and no APs (Hardie and Spruston, 2009). We tested this idea in the apical tree of the L5 pyramidal neuron (Fig 6A). If two branches receive different synaptic activation (quantified as the product between local impedance and synaptic conductance, see methods), the voltage difference between these branches will be larger when they are separated by a higher *I_Z_* (Fig 6C), and therefore the probability increases to robustly potentiate the preferred branch while the non-preferred branch is depressed. For the selected branches (Fig 6A), *I_Z_* originally was too low (Fig 6D, G), and thus the evolution of synaptic weights in the preferred and non-preferred branches was positively correlated (Fig 6F). Activation of inhibitory synapses near the bifurcation point increased *I_Z_* (Fig 6B, black vs blue tree), thus anti-correlating the weight evolution (Fig 6F) and enabling branch-specific learning (Fig 6E, G).

**Figure 6:**
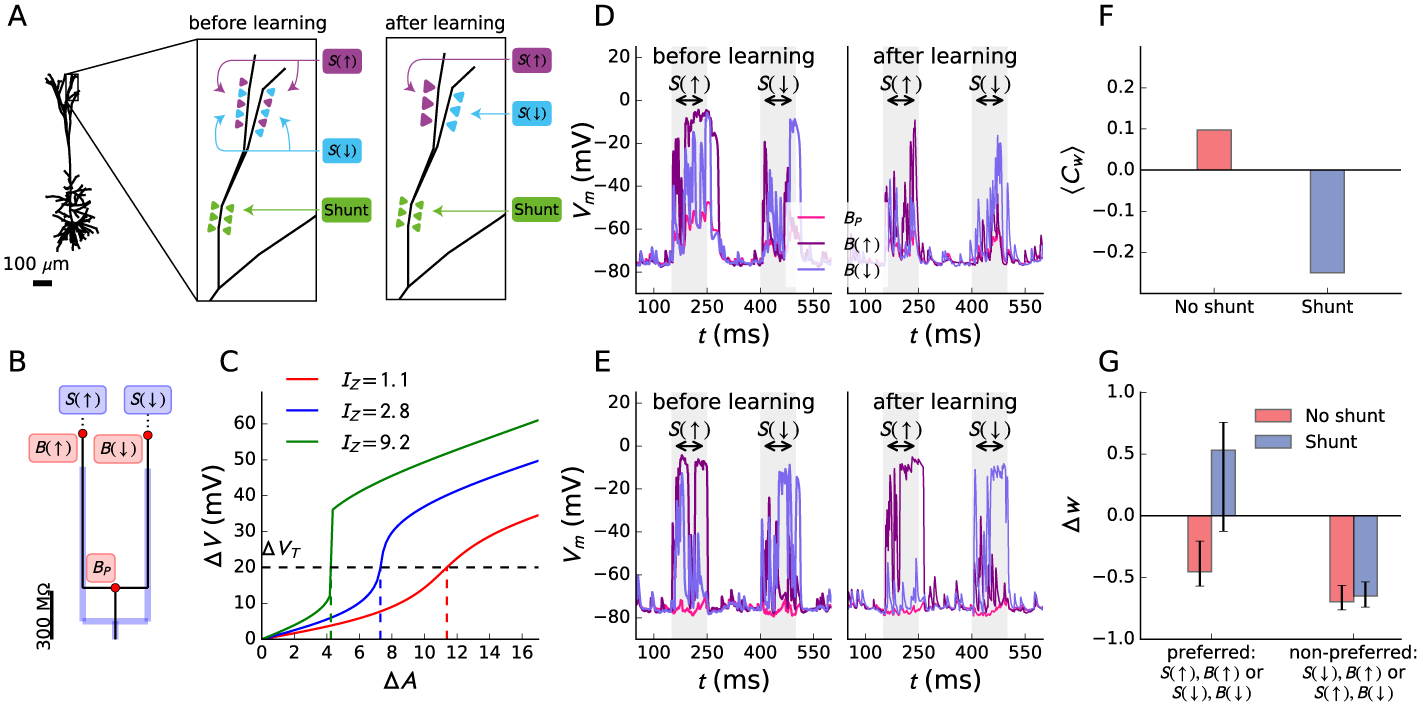
Enabling branch-specific learning with shunting conductances. **A:** Sketch of the situatio**n:** before learning, two sibling branches are both targeted by synapses coding for either an up stimulus *S*(↑) (purple) or a down stimulus *S*(↓) (blue). One branch *B*(↑) receives more synapses of the former stimulus and the other branch *B*(↓) more of the latter stimulus. The branches should learn to be sensitive only to the stimulus that was initially the most prevalent (the ‘preferred’ stimulus). The shunting inhibition in the parent branch *B_P_* had a time-averaged conductance of 12 nS when active. **B:** Schematic of the NET for the depicted situation. **C:** The difference in voltage across sister branches Δ*V* increases as a function of the difference in activation Δ*A*. For higher *I_Z_*, this increase is steeper. Consequently, a given threshold Δ*V* (black dashed line) is reached for lower Δ*A* (colored vertical lines). D, **E:** The learning task without (D, effective *I_Z_* =3.0) or with (E, effective *I_Z_* =8.5) shunting inhibition. Each epoch, both stimuli are presented for 100 ms, with 150 ms intervals in between them, for a total of 20 epochs. The initial and final epochs are shown. **F:** Correlation between the average weights of the synapses in the preferred and non-preferred branches during stimulus presentation, averaged over all epochs and 20 repetitions of the learning task. **G:** Bar plot of the weight difference after the final epoch for all repetitions of the learning task. The bar lengths denote the medians of the weight difference distributions and the error bars the 25–75 percentiles. With shunt, the neuron is able to successfully learn the task.

## Discussion

In this work we have formalized dendritic compartmentalization in three steps. First, recognizing that independence across dendritic sites increases continuously as a function of their electrical separation, we have introduced the impedance based independence index (*I_Z_*) and have shown that this single number suffices to quantify independence between sites. Second, we have shown that for *I_Z_* ≳ 10 pairs of dendritic sites are truly independent when they receive independent inputs. Third, we have employed the NET to design an algorithm that, given a threshold *I_Z_*, yields the maximal number of sites that can coexist on the dendritic three and for which each pair is separated by an *I_Z_* larger than or equal to the threshold value.

We have then performed this analysis on a number of cell classes (Fig S2) and found, to our surprise, that many branches are not separated by *I_Z_* values required for independence. Impedance drops across dendritic bifurcations lead to large voltage attenuation (Fig 3), and studying only simple input scenarios may lead to the idea that sibling branches automatically constitute independent subunits. We nevertheless found that in more realistic input scenarios, when both branches receive an ongoing barrage of inputs, relatively small transfer impedance values still lead to a large degree of cooperativity between synapses (Fig 4). Only for relatively large separations (*I_Z_* ≳ 10), branches are able to truly function independently.

The heterogeneity of brain states (Wilson and Kawaguchi, 1996) as well as the observation that the electrical length of nerve fibers changes with the amount of background conductance (Segev, 2000) led us to explore the possibility that the NET, and hence independence, could be modified dynamically by spatiotemporal input patterns. We found that up-states (Rudolph and Destexhe, 2003; Destexhe et al., 2007), which increase membrane conductance globally across the dendrite, tend to reduce the impedance associated with the root node of the NET more strongly than impedances associated with leaf nodes. Hence, during up states, independence across branches increases (Fig 5A-D). We also found that shunting inhibition can increase independence in a highly localized fashion, thus resulting in the precise tuning of compartmentalization (Fig 5E-H).

How many branches on the dendritic tree lie within in the useful range for dynamic recompartmentalization? In pyramidal cells, compartments numbers double for values of *I_Z_* between 3 and 10 and in smaller cells these numbers increase five-fold (Fig S2). This suggests that global conductance input increases the number of compartments two-to five-fold, depending on the cell’s properties. Inspecting pairwise independence, we determine that in pyramidal cells 5% to 10% of terminal pairs are separated by *I_Z_* values between 3 and 10 (mainly terminals on the same main branches). In smaller cells, up to 60% of pairs fall within these values. On average, these pairs are made independent by shunting conductances of 5 to 15 nS (Fig S2).

To explore the functional consequences of dynamic dendritic compartmentalization, we equipped synapses on dendritic sibling branches with a voltage-based plasticity rule (Clopath et al., 2010) and explored whether these synapses could learn independently. Consistent with the aforementioned observations, we found that a value of *I_Z_* ≃ 3 was too small for these branches to learn independently. Nevertheless, shunting inhibition that increased *I_Z_* to 8.5 allowed the synaptic weights in both branches to evolve in an independent fashion. Interestingly, recent experimental data suggest that such phenomena may occur in-vivo (Cichon and Gan, 2015). These results also suggest that the computation a neuron is engaged in may vary across brain states: when background conductance is high, neurons may engage primarily in local dendritic learning, whereas otherwise they may favour associative output generation. Varying phases of neuronal processing are hypothesized to occur in theories of how networks of neurons learn (Guergiuev et al., 2016).

Across the brain, neurons take on a wide variety of dendritic morphologies. We have shown here for the first time how these dendritic trees compartmentalize at rest and during dynamic input regimes. The behavioural relevance of up states (Destexhe et al., 2007), the specificity of inhibitory targeting (Bloss et al., 2016) and the observation that abolishing somatostatin-positive interneuron activity diminishes branch-specific learning and motor task performance (Cichon and Gan, 2015) suggest that dynamic compartmentalization is ubiquitous in normal brain function, with far-reaching consequences for memory formation (Kastellakis et al., 2016) and capacity (Poirazi and Mel, 2001; Wu and Mel, 2009).

Because the NET provides a general framework for understanding synaptic interactions, it allows for easy integration of previously described synaptic coupling mechanisms (Larkum et al., 1999) and may improve our understanding of the computational role of spatial synapse distributions (Gökçe et al., 2016). Furthermore, we have devised an efficient inversion algorithm (see extended data), so that a performant NET simulation paradigm can be designed. This will enable scientists (i) to implement efficient dendrite models for network simulations and (ii) to create abstract dendrite models that implement dendritic computations in a minimal fashion. Taken together, the NET can be seen as a computational description of the morphological neuron, complementary to the well-known biophysical description, and the algorithm to derive it as a translation from biophysics to computation.

## Methods

### Biophysical modelling

#### Morpohologies

Three exemplar morphologies were used for the analysis: a cortical stellate cell (Wang et al., 2002) (Fig 2A), a hippocampal granule cell (Carim-Todd et al., 2009) (Fig 2B) and a cortical pyramidal cell (Hay et al., 2011) (Fig 2C). These morphologies were retrieved from the NeuroMorpho.org repository (Ascoli, 2006), except the pyramidal cell, which was retrieved from the ModelDB repository (Hines et al., 2004). Cell morphologies used in our wider cortical analysis were retrieved from the Blue Brain Project database (Markram et al., 2015).

#### Physiological parameters

Physiological parameters for the morphologies were set according to Major et al. (Major et al., 2008): the equilibrium potential was −75 mV, the membrane conductance 100 *µ*S*/*cm^2^, the capacitance 0.8 *µ*F*/*cm^2^ and the intracellular resistance 100 Ω · cm.

To generate somatic APs, we used the fast inactivating Na^+^ current (*g*Nat) and the fast, non-inactivating K^+^ current (*g*Kv3.1) previously employed in cortical models (Hay et al., 2011). Channel densities were: 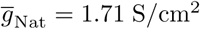 and 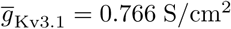. The leak current was then fitted to yield a membrane time scale of 10 ms and an equilibrium potential of −75 mV.

AMPA and GABA synaptic input currents were modeled as the product of a conductance profile, given by a double exponential shape (Rotter and Diesmann, 1999), with a driving force (Jack et al., 1975):

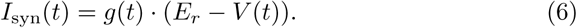

For AMPA synapses, we used rise resp. decay times *τ_r_* =0.2 ms, *τ_d_* = 3 ms for the conductance window and a reversal potential *E_r_* = 0 mV, while for GABA synapses we used *τ_r_* =0.2 ms, *τ_d_* = 10 ms and *E_r_* = −80 mV. For N-methyl-Daspartate (NMDA) channels (Jahr and Stevens, 1990), the synaptic current had the form

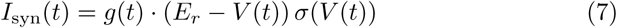

and the rise resp. decay time were *τ_r_* =0.2 ms, *τ_d_* = 43 ms, and *E_r_* = 0 mV, while *σ*(*V*) was the sigmoidal function employed by (Behabadi and Mel, 2014) to model the channels’ magnesium block: 1

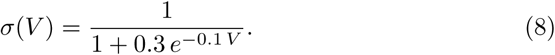

In the remainder of this work, we will refer to the voltage-dependent factors in the synaptic input current as the ‘synaptic voltage dependence’ (SVD), denoted by *f*(*V*). Hence, for AMPA or GABA synsapses

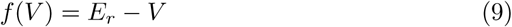

and for NMDA synapses

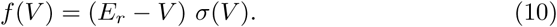

Note that when we refer to the conductance of a simple synapse, we mean the maximum value *g*_max_ of its conductance window. For a synapse that has AMPA and NMDA components (to which we will simply refer as an NMDA synapse), the conductance is the maximal value of the AMPA conductance window, and the conductance of the NMDA component is determined by multiplying the AMPA conductance value with an NMDA ratio *R*_NMDA_, that was set to be either 2 or 3.

#### Plasticity

In our simulations with plasticity, we use a voltage dependent spike timing dependent plasticity rule (Clopath et al., 2010; Bono and Clopath, 2017) where the evolution of the weight *w*(*t*) of a given synapse depends both on the postsynaptic voltage and the presynaptic AP inputs. This leads to a synaptic current of the following form:

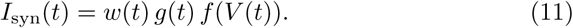

In all our simulations the initial weight *w*(*t* = 0) was 1 and, during the simulation, the weight could fluctuate in the interval [0, 2].

#### Compartmental models

To construct and simulate compartmental models of the cells, we used the neuron simulator (Carnevale and Hines, 2004). Compartment sizes were set to be smaller than or equal to the size given by the lambda rule (Carnevale and Hines, 2004).

#### Green’s function and the separation of variables

To derive NETs, we rely on the Green’s function (GF) (Koch, 1998; Wybo et al., 2013, 2015) *Z*(*x*, *x*′, *t*). The GF is a function of three variables: two locations *x* and *x*′ along the dendritic arborization and a temporal variable *t*. We compute the GF in an exponential basis:

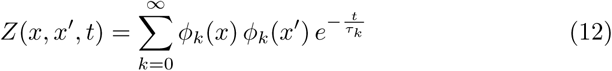

by using the separation of variables (SOV) method (Major et al., 1993; Major and Evans, 1994). Note that it is a property of the cable equation that the GF is symmetric in the spatial coordinates (Koch, 1998), so that *Z*(*x*,*x*′,*t*) = *Z*(*x*′, *x*,*t*). Usually, a fixed set of discrete locations relevant for the problem at hand is chosen on the neuron. Hence, the GF only needs to be evaluated at these locations, and a discrete set of temporal *kernels* is obtained. A member of this set will be denoted as *Z_x_*, *_x_*_′_ (*t*), to highlight the difference between the now discrete indices *x* and *x*′ and the continuous variable *t*.

To compute the output voltage *V_x_*(*t*) at location *x* for a given input current *I_x_*_′_ (*t*) at location *x*′, one needs to compute the convolution of the GF evaluated at *x* and *x*′ with this input current (Koch, 1998):

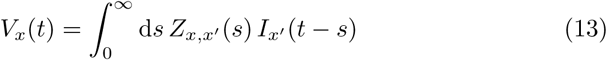

for which we will use the shorthand

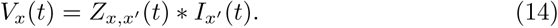

Since *Z_x, x_*_′_(*t*) converts current into voltage, we will refer to it as an ‘impedance kernel.’ The total surface under the impedance kernel is the steady state impedance:

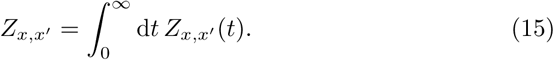

In the rest of the text, it will be understood that *Z_x, x_*_′_ without temporal coordinate refers to the steady state impedance – which we will simply call ‘the impedance’ for brevity – while *Z_x, x_*_′_(*t*) is the temporal impedance kernel. To unclutter the notations, we will not make this distinction for other variables; the temporal dependence will be omitted by default. Following this convention, equation (14) will be written as *V_x_* = *Z_x, x_*_′_(*t*) ∗ *I_x_*_′_, where it is implied that both *I_x_*_′_ and *V_x_* are time dependent quantities since *Z_x, x_*_′_(*t*) is the temporal impedance kernel. Conversely, writing *V_x_* = *Z_x, x_*_′_ *I_x_*_′_ means that *Z_x, x_*_′_ is the steady state impedance value, and thus *I_x_*_′_ and *V_x_* will be steady state values too. Note that currents in this text will be expressed in nano ampere (nA) and voltages in milli volt (mV). Consequently, impedances will be in mega ohm (MΩ).

#### Synaptic activation

The eventual steady state voltage *V_x_* obtained after activating a synaptic conductance at location *x* depends for a large part on the input impedance *Z_x,x_*. Following (13), it can be obtained as a solution of the equation

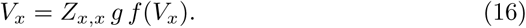

This solution *D* is thus a function of the product of impedance and conductance:

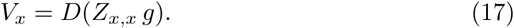

We refer to this product as the synaptic activation *A_x_* = *Z_x,x_ g* and note that it is a dimensionless quantity. Consequently, it is a convenient quantity that does not depend on local morphological constraints to determine whether an input will be strong enough to reach a certain voltage threshold, for instance to elicit an NMDA spike or to potentiate a synapse.

### Neural Evaluation tree

#### Mathematical formulation

In the NET framework, the voltage at a node can be expressed as:

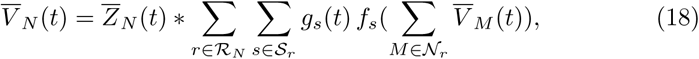

where 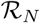 is the set of all input regions a node *N* integrates, 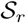 the set of all synapse types at region *r* (with *g_s_* their conductance and *f_s_*(·) their SVD) and 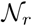 the set of all nodes that integrate region *r*. 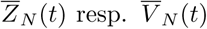 denote the impedance kernel resp. voltage at node N. Note that at the NET leafs, each node integrates inputs from only one region. Note furthermore that the matrix associated with this system can be inverted in *O*(*n*) steps (with *n* the number of nodes), so that efficient NET simulations can be designed (see supplementary information).

#### Derivation

To derive the NET, we order all locations along the dendritic arborization in a depth-first manner (Russell and Norvig, 2003), so that the impedance matrix (Cuntz et al., 2010) has a highly organized structure (Fig 1C, D). Generally an even blue surface covers most of the matrix, representing the transfer impedances between the main dendritic branches. There are also smaller square regions of light blue or green closer to the diagonal, representing sibling branches that are electrically closer to each other than to different main branches. Finally, the small squares along the diagonal coloured yellow and red are the thin dendritic tips with high input impedances. Consequently, dendritic tips that lie within the same light blue or green square are closer together electrotonically than tips within different squares, as the transfer impedances connecting them are much higher. A NET tree graph structure hence imposes itself naturally: the dendritic tips constitute the leafs of the tree, their parent node combines multiple adjacent tips and the root node in turn binds all these nodes together. To derive the NET tree graph, we define an impedance step Δ*Z* and execute the following recursively:

1. Let *k* denote the step number. We assume that a value *Z*_min_,*_k_* is at our disposition. In the first step this value is 0, in later steps it is given by the previous steps. A value *Z*_max_,*_k_* is also determined, in the first step as the somatic input impedance (*Z*_00_ with our depth-first ordering of the impedance matrix) and in later steps by *Z*_max_,*_k_* = *Z*_min_,*_k_* +Δ*Z*.
2. The kernel of the current node is constructed as the average of all impedance kernels associated with points in the impedance matrix for which *Z*_min_,*_k_* ≤ *Z_ij_* <*Z*_max_,*_k_* (coloured blue in Fig 1E). This approach is justified, as impedance kernels of similar magnitude have similar time scales (Fig 1B). Then, the kernels of all underlying nodes are subtracted from this average kernel.
3. The next nodes are determined by looking at the input impedance, located on the diagonal of the impedance matrix. Due to the depth-first ordering, new nodes can be identified as uninterrupted intervals on this diagonal where *Z_ii_* >*Z*_max_,*_k_* (Fig 1F). For each of these intervals, a new child node is constructed by repeating step 1 with the impedance matrix restricted to the interval (indicated in Fig 1G by the red squares) and with *Z*_min_,*_k_*+_1_ = *Z*_max_,*_k_*.

Generally, one is only interested in a subset of input regions on the dendrites, so that the NET can be pruned of nodes that do not integrate inputs. Furthermore, if multiple nodes integrate the same input region, they can be replaced by one node whose impedance kernel is the sum of all impedance kernels of the original nodes.

The average error of this approximation depends on Δ*Z*, and approaches its minimum value for Δ*Z* ≲ 20 MΩ (Fig S3A).

#### Distal and proximal regions

The aforementioned algorithm successfully constructs NETs of dendritic arborization where the variations in transfer impedance between branches are small compared to their average values (Fig S3B-D). If these variations are large, as is the case in pyramidal cells (Fig S3D), interactions between proximal and distal dendritic domains are overestimated. In pyramidal cells, distal domains are only connected to the soma by one or a few large dendritic branches. Hence, there are many points of low soma to dendrite impedance *Z*_0_*_x_*, many of high *Z*_0_*_x_*, and relatively few of intermediate *Z*_0_*_x_*. By consequence, the histogram of all dendrite to soma transfer impedances has two main modes, whose boundary (Delon and Desolneux, 2007) indicates the domains. Any region with *Z*_0_*_x_* above this boundary will belong to the proximal domain (node *c*_0_ in Fig S3D), whereas connected regions with *Z*_0_*_x_* below this boundary will constitute distal domains (coloured red in Fig S3D, node *c*_1_). Algorithmically, we determine the kernel of the root node as the average of all transfer impedance kernels between proximal and distal branches. Then, we start the recursive procedure as before, but with the impedance matrices restricted to the different domains *c_i_*.

#### Predicting spikes: linear terms

The effective NET transfer impedance between dendrite and soma is either constant or can take on a proximal and a distal value. Spike prediction is refined further by defining linear terms that capture the precise transfer impedance between input regions and soma. These kernels only contribute to the voltage at the soma, and thus have no influence on the intra-dendritic synaptic interactions. Mathematically, their contribution to the somatic voltage can be written as:

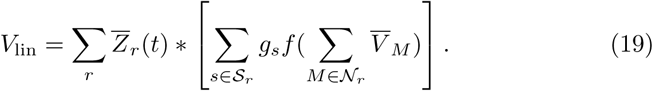

### Independence and compartmentalization

#### True independence

The leafs of the NET only receive inputs from a single region. Nevertheless, they are not per se independent from the other synapses, since the SVD in their input current still depends on all nodal voltages on the path to the root:

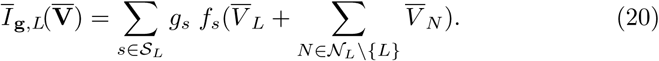

This current will become truly independent from all other synapses if

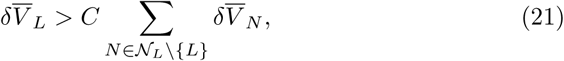

with *C* a (large) number that has to be determined empirically and 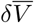 denoting the short term fluctuations of 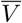 around a long term average 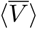. Here, short term means the time-scale on which neurons convert electrical inputs to output. As NMDA synapses have a decay time constant of 43 ms, this averaging time-scale should be at least somewhat larger than the NMDA time constant (we chose ~ 200 ms). Then (22) can be approximated as follows:

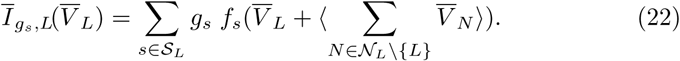

The long term average in this equation is only influenced very little by the instantaneous values of the synaptic conductances and can hence be seen as a constant, i.e. a fixed parameter in the equation:

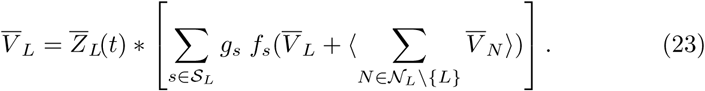

Consequently, the solution for 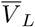 will not depend on the instantaneous values of the synaptic conductances at other locations.

#### Estimating independence

Whether condition (21) holds depends on the structure of the NET as well as the relative size of the synaptic inputs. We assume that synaptic conductances in physiological regimes are of similar magnitude. In this case, condition (21) becomes a condition on the impedances:

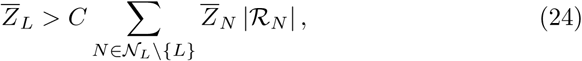

where 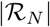 denotes the number of regions node *N* integrates. When we are interested in determining whether a pair of regions *r_i_* (integrated by the leafs *L_i_*, *i* =1, 2) can act independently, we can consider a reduced tree with two leafs, obtained by pruning all nodes associated with other regions. The new tree then has leaf impedances 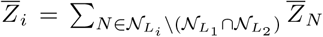 (i.e. a sum over impedances of nodes that integrate one region but not the other) and a root impedance 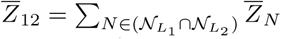 (i.e. a sum over impedances of nodes that integrate both regions). Then, regions *r_i_* are independent if:

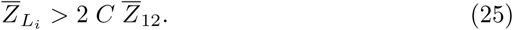

For mutual independence between *r*_1_ and *r*_2_, this equation has to hold for both *i* = 1 and *i* = 2. To summarize these two conditions in a single expression, we defined the ‘impedance-inferred independence index’ (*I_Z_*):

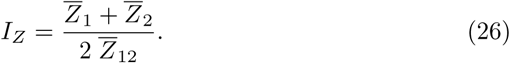

Then, if (25) holds for both regions, the following condition also holds:

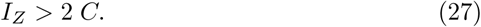

Note that this is a necessary, but not a sufficient condition for mutual independence. However, as shown throughout the main manuscript, when asymmetry is not too high *I_Z_* is, despite its simplicity, a surprisingly accurate measure.

#### Compartmentalization

Previously we discussed the conditions under which a single input site can be considered independent from the rest of the input regions. Nevertheless, when inputs are distributed in an almost continuous fashion along the dendritic arborization, such sites may not exist. It can be expected, however, that the structure of the dendritic tree favours a grouping of inputs, such that inputs belonging to different groups are all mutually independent but inputs belonging to the same group are not. A grouping of this type for homogeneously distributed inputs along the dendritic arborization, and where inputs belonging to different groups have an *I_Z_* above a certain threshold, will be called a compartmentalization of that dendritic tree for that given *I_Z_*. Note that in such a compartmentalization, *not all input sites can belong to a group*, as there will have to be at least some space between compartments.

How can such a compartmentalization be found? First, we remark that there is no unique answer to this question. Consider a forked dendritic tip. It may happen that inputs within each sister branch are independent from the rest of the dendritic tree, but the branches are not independent from each other. Furthermore, because of a steep impedance gradient within the branch, inputs at the bifurcation point may not be independent from the rest of the tree. Because of the first constraint, both tips can not form separate compartments, whereas because of the second constraint, they can not be grouped into a single compartment either. Hence only one branch can be chosen, and either choice forms a valid compartmentalization.

We implemented an algorithm that proposes, using the NET and given an *I_Z_*, a compartmentalization that maximizes the number of compartments. We note that if a node *N* in the NET tree forms a valid compartment, all nodes in the subtree of *N* are part of the same compartment, since their *I_Z_* to other compartments will be higher than the *I_Z_* of *N*. Hence, our algorithm will simply return a set of nodes, where it is understood that a compartment associated with a node from this set is its whole subtree. Our algorithm proceeds in three steps:

1. We determine a ‘tentative’ compartmentalization. For each node *N* in the NET tree, we examine the bifurcation nodes *B* on the path 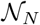 from *N* to the root. We check whether the following condition holds

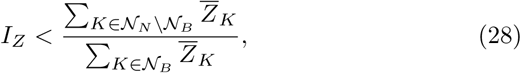

with 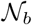 the path from *B* to the root. If this condition is true for two nodes *N* and *M* that have *B* on their respective paths to the root, and where 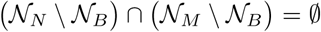, these nodes will be separated by at least the required *I_Z_*. Hence, we say that *N* is a tentative compartment with respect to *B*.
2. In a second step, we remove all leafs from the tree that could not possibly be separate compartments. To do so, we look at the highest order bifurcation *B* and its child leafs. Then, if at least two child leafs are tentative compartments with respect to *B*, the other leafs are removed. Otherwise, all child leafs but the one with largest impedance are removed. Note that in the latter case, *B* is not a bifurcation anymore and consequently will not induce tentative compartments. We continue to cycle through the bifurcation nodes of highest order until no more nodes can be removed.
3. In a final step we assign the compartments. As we are now sure that every leaf is part of a separate compartment, we start at the leaf, find the nearest bifurcation node in the NET tree, and then recursively find the lowest order node that is still a tentative compartment of *B*. This node will be a compartment node in the final compartmentalization.

#### Simulation-specific parameters

**Parameters figure 1**. We evaluated the impedance matrix in panel B at 10 *µ*m intervals. In the simulation depicted in panels J and K, the synapses contained only an AMPA component with *g*_max_ = 10 nS and no active channels were inserted in the soma.

**Parameters figure 2**. 100 NMDA (*g*_max_ = 4 nS and *R*_NMDA_ = 3) and 100 GABA (*g*_max_ = 2 nS) synapses were inserted on the morphology and activated with Poisson spike trains of 1 Hz.

**Parameters figure 3**. The synapse at region 1 was an NMDA synapse with *g*_max_ = 1 nS and *R*_NMDA_ = 3. The synapse received 5 input spikes in a 2.5 ms interval in order to trigger an NMDA spike

**Parameters figure 4**. For the simulations in panels C-D, NMDA synapses (*R*_NMDA_ = 3) were used. Their *g*_max_ and incoming Poisson rate were optimized to utilize the full range of the NMDA non-linearity. For the simulations in panel E, NMDA (*g*_max_ = 2 nS and *R*_NMDA_ = 2) and GABA (*g*_max_ = 1 nS) synapses were used and their rates were also optimized to utilize the full range of the NMDA non-linearity.

**Parameters figure 5**. For the simulations in panel C, the main synapses contained only an AMPA component with *g*_max_ = 5 nS. To simulate the high-conductance state, 200 AMPA and 200 GABA synapses (*g*_max_ =0.5 nS) were distributed evenly across the neuron. Each AMPA synapse was stimulated with a Poisson spike train of 5 Hz. The rate of stimulation for the GABA synapses was tuned to achieve a balanced input. To recompute the tree structures for panels B and D, the time-averaged conductances of all background synapses were inserted in the morphology as static shunts.

The inhibitory synapse in panels E-H had *g*_max_ =2.3 nS and was activated at a steady rate of 200 Hz, so that it’s total time-averaged conductance was around 5 nS. For the simulations in panel H, we inserted NMDA synapses (*R*_NMDA_ = 3) in both branches and stimulated them with a rate of 200 Hz. Note that these inputs could come from multiple presynaptic cells. Due to the linearity of the conductance dynamics however all spikes can be taken to add to the same conductance and can hence be modeled as a single synapse. The maximal conductance *g*_max_ of the NMDA synapse was optimized to obtain an average depolarization of −40 ± 2.5 mV in each branch, a target value which yields parameters that allow exploitation of the full range of the NMDA non-linearity.

**Parameters figure 6**. In both simulations with and without shunting inhibition, noise was implemented at all three locations using AMPA (*g*_max_ =0.1 nS) and GABA (*g*_max_ =0.2 nS) synapses. Both were stimulated with Poisson spike trains of resp. 33 Hz and 83.1 Hz (tuned to achieve balance). The shunting inhibition in the parent branch was implemented by a GABA synapse (*g*_max_ = 2 nS) receiving a Poisson train with a rate of 277 Hz (tuned to reach a time-averaged conductance of 12 nS) during the 100 ms learning intervals. Note that this single conductance could again represent multiple synapses.

Stimulus-specific innervation patterns were: *S*(↑) to *B*(↑): 5 synapses, *S*(↑) to *B*(↓): 2 synapses, *S*(↓) to *B*(↑): 2 synapses and *S*(↓) to *B*(↓): 5 synapses. These synapses were all NMDA synapses (*g*_max_ =0.6 nS, *R*_NMDA_ = 2) that where activated in the learning intervals with Poisson trains at a rate of 33.3 Hz without the shunting inhibition and 39.8 Hz with the shunting inhibition (to compensate for the loss in input impedance in both branches).

**Parameters supplementary information figure S1**. In panel D, synapse 1 was a non-plastic NMDA synapse (*g*_max_ = 1 nS, *R*_NMDA_ = 3) and synapse 2 a plastic synapse with the same parameters. When on, synapse 1 received a tonic spike train with a rate of 113 Hz (to yield a time-averaged activation *A*_1_ ≃ 15). Synapse 2 received rates ranging from 0 to 113 Hz, corresponding to the data points at different activations in the figure in the figure. For the simulations in panel H, we inserted NMDA synapses (*R*_NMDA_ = 3) in both branches and stimulated them with a rate of 200 Hz. Their maximal conductance *g*_max_ was optimized to obtain an average depolarization of −40 ± 2.5 mV in each branch.

## Supplementary information

Figures S1-S3, description of NET matrix inversion algorithm and analytical results for conductance inputs

## Acknowledgments

WW and MOG were supported by funding from the ETH Domain for the Blue Brain Project (BBP) and the European Union Seventh Framework Program (FP7/2007–2013) under grant agreements no. FP7–269921 (BrainScaleS), FP7-604102 (THE HUMAN BRAIN PROJECT), as well as EU grant agreement no. 720270 (HBP SGA1). WW and BTN designed the research. WW performed the mathematical modelling, implemented the models and ran the simulations. WW, BTN and MOG analysed the results and wrote the paper. We thank Prof. Idan Segev and Prof. Wulfram Gerstner for helpful advice and discussions and Dr. Luc Guyot and Dr. Aditya Gilra for proofreading the manuscript.

